# Accelerating *de novo* SINE annotation in plant and animal genomes

**DOI:** 10.1101/2024.03.01.582874

**Authors:** Herui Liao, Yanni Sun, Shujun Ou

**Affiliations:** Department of Electrical Engineering, City University of Hong Kong, Hong Kong SAR, China; Department of Molecular Genetics, The Ohio State University, Columbus, OH, 43210, USA

**Keywords:** Genome annotation, Transposable element, SINE identification

## Abstract

Genome annotation is an important but challenging task. Accurate identification of short interspersed nuclear elements (SINEs) is particularly difficult due to their lack of highly conserved sequences. AnnoSINE is state-of-the-art software for annotating SINEs in plant genomes, but its homology-based module is not available for animals and it is computationally inefficient for large genomes. Therefore, we propose AnnoSINE_v2, which extends accurate SINE annotation for animal genomes with greatly optimized computational efficiency. Our results show that AnnoSINE_v2’s annotation of SINEs has over 20% higher F1-score compared to the existing tools on animal genomes and enables the processing of complicated genomes, like human and zebrafish, which were beyond the capabilities of AnnoSINE_v1. AnnoSINE_v2 is freely available on Conda and GitHub: https://github.com/liaoherui/AnnoSINE_v2.

## Introduction

Short interspersed nuclear elements (SINEs) are interspersed transposable elements in eukaryotic genomes, playing key roles in enhancing genome complexity, regulating gene expressions, and generating novel genes [1,2]. Accurate annotation of genome-scale SINEs is essential for various downstream analyses, including comparative genomics and evolutionary studies [3]. However, the short length and lack of well-conserved sequences in SINEs pose challenges for existing annotation tools, leading to low sensitivity or high false positive rates [4]. To address these limitations, AnnoSINE, a recently developed SINE annotation tool, employs a hybrid approach that combines homology-based and *de novo* search strategies. By integrating a comprehensive set of sequence features, AnnoSINE effectively identifies SINE candidates while mitigating false discoveries. While AnnoSINE has improved SINE annotation in plants, its homology-based annotations cannot be applied to animal genomes due to the lack of animal-origin profile hidden Markov models (pHMMs). Furthermore, it is computationally prohibitive for AnnoSINE to annotate large genomes due to algorithmic inefficiency. In this study, we introduce AnnoSINE_v2, an update that enables SINE annotation in animal genomes with notable efficiency improvements achieved through multi-threading and optimization strategies.

## Results

To enable homology-based SINE identification in animal genomes, we collected 118 SINE families represented by pHMMs from the DFAM database (Supplementary Table S1) [5]. These animal pHMMs effectively extend AnnoSINE_v2’s ability to annotate SINEs within animal genomes. To improve the computational efficiency, we have optimized various time-consuming functions within the original pipeline and incorporated the utilization of multithreading. To assess the performance of AnnoSINE_v2 with the original AnnoSINE (referred to as “AnnoSINE_v1”), we benchmarked AnnoSINE_v2 with AnnoSINE_v1 using rice and *Arabidopsis thaliana* genomes. The results indicate that AnnoSINE_v2 achieves comparable performance to AnnoSINE_v1 (Supplementary Table S2).

To evaluate AnnoSINE_v2’s performance in animal genomes, we conducted a benchmark experiment using zebrafish, mouse, and human genomes with curated SINE annotations downloaded from the UCSC genome browser [6]. According to [6], half of annotations are computed internally at UCSC, while the remaining annotations are contributed by research scientists from around the world. In the subsequent analysis, these annotations were served as the ground truth. In this evaluation, we also included two popular SINE annotation tools, SINE-Finder [7] and SINE-Scan [8], for comparison. As shown in Fig. 1, AnnoSINE_v2 exhibits an overall superior performance to SINE-Finder and SINE-Scan across all tested genomes. Notably, the specificity of AnnoSINE_v2 exceeds 90% in all three genomes. AnnoSINE_v2 also achieves higher F1 scores than SINE-Finder and SINE-Scan, with increases of 22%∼25% observed in zebrafish, mouse, and human genomes. Although SINE-Finder and SINE-Scan exhibit higher sensitivity in mouse and human genomes, they also have ≥20% higher false positive rates (Fig. 1). These data indicate that AnnoSINE_v2 identifies substantially fewer false predictions in the tested animal genomes.

**Fig. 1.**
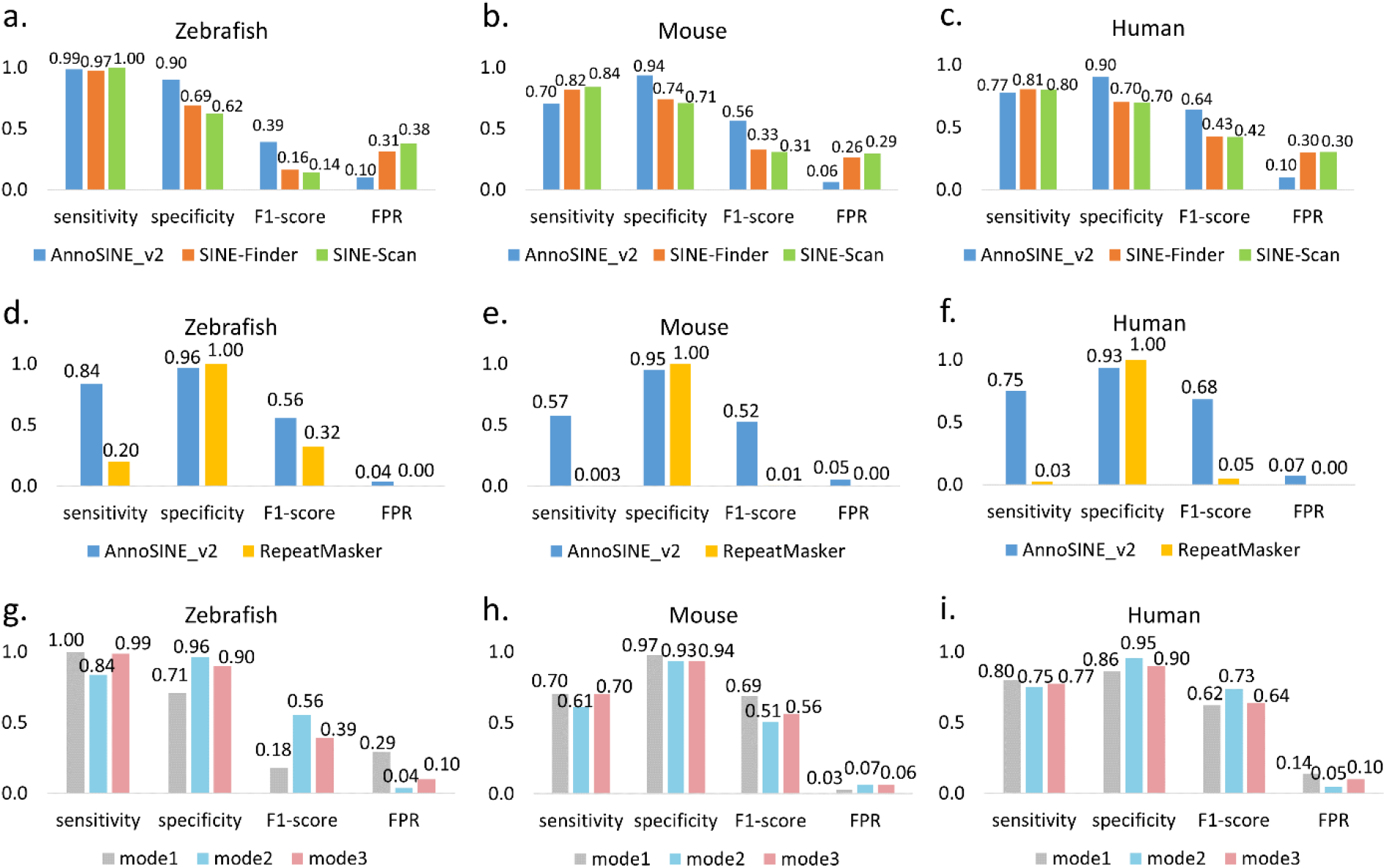
(a-f): The performance of AnnoSINE_v2, SINE-Finder, SINE-Scan, and RepeatMasker in annotating SINEs across three animal genomes (zebrafish, mouse, and human). (g-i): The performance of AnnoSINE_v2 under different modes. FPR: false positive rate. Evaluations were based on whole-genome SINE annotations and their total length in bp. When benchmarking AnnoSINE_v2 and RepeatMasker, all SINEs from zebrafish, mouse, and human genomes and closely related species were removed from the database to test their *de novo* SINE identification ability.

Since AnnoSINE_v2 also recognizes SINE elements based on HMM profiles of existing SINE families, having such families included in the testing library will inflate the sensitivity results. To further access the performance of AnnoSINE_v2 in *de novo* SINE annotation, we eliminated all SINEs from the zebrafish, mouse, and human genomes available in the DFAM database. Then, we utilized this modified database to benchmark the tools in the aforementioned genomes. In addition, we employed the remaining SINE HMM profiles as the custom library for RepeatMasker, enabling a comparison of its performance with AnnoSINE_v2. AnnoSINE_v2 exhibited more competitive performance with increased sensitivity compared to RepeatMasker, highlighting its enhanced ability for *de novo* SINE identification (Fig. 1). Furthermore, to assess the impact of the homology-based method versus the structure-based method, we also investigated the performance of AnnoSINE_v2 under different modes (Fig. 1 g-i). As shown in the figure, mode 1 typically achieves higher sensitivity, while mode 2 tends to exhibit higher specificity. In addition, neither mode 1 nor mode 2 can maintain the best performance in all tested species. In contrast, mode 3 consistently achieves the second-best F1-score, indicating its robust performance.

To investigate the influence of parameters, we compared AnnoSINE_v2 and SINE-Finder on the rice genome using different parameter settings. AnnoSINE_v2 provides parameters such as “-s” (maximum threshold of SINE boundary shift) and “-minc” (minimum threshold of copy number for each element), while SINE-Finder offers parameters like “-t” (target site duplications (TSD) mismatch tolerance) and “-s” (TSD score cutoff). By adjusting these parameters, we assessed their impact on performance (Fig. 2 and Supplementary Table S3). We found that increasing the value of “-s” in AnnoSINE_v2 improved the true positive rate (TPR) but also resulted in a higher false positive rate (FPR), whereas increasing “-minc” reduced both FPR and TPR. Similarly, for SINE-Finder, increasing “-t” enhanced TPR but also increased FPR, while increasing “-s” reduced both TPR and FPR. These results indicate that adjusting parameters such as SINE boundary shift or TSD score cutoff can influence the tradeoff between sensitivity and specificity of these two tools. Notably, AnnoSINE_v2 consistently maintained a low FPR across parameter variations, whereas SINE-Finder exhibited significant variability in response to parameter changes.

**Fig. 2.**
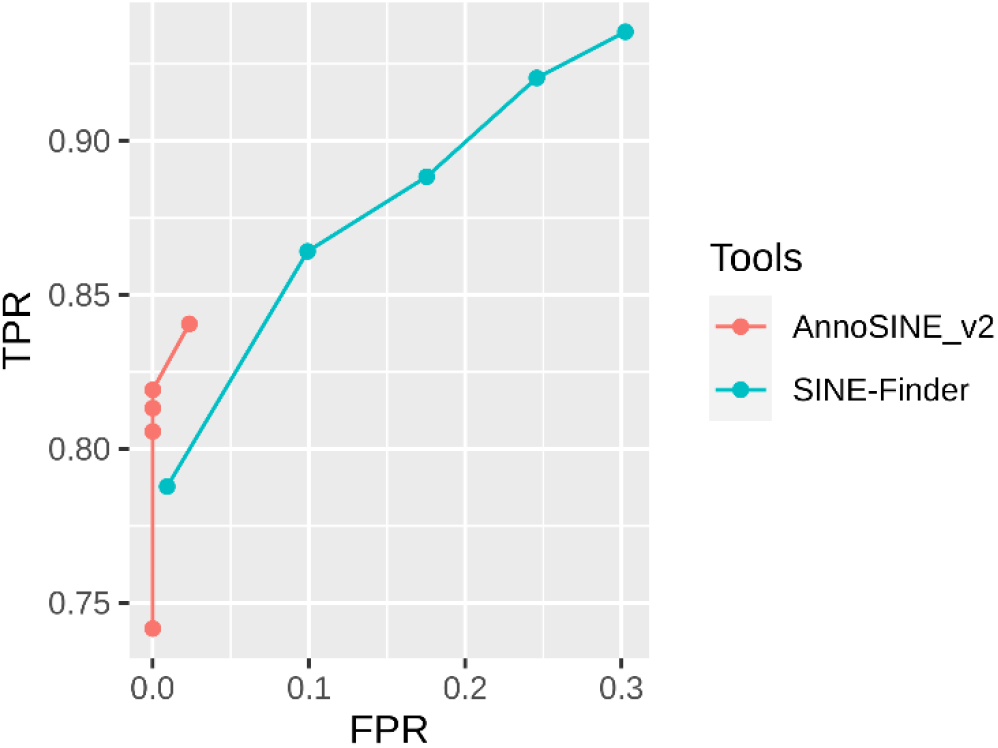
The ROC curve of AnnoSINE_v2 and SINE-Finder under different parameter settings, including “-s” (maximum threshold of SINE boundary shift) and “-minc” (minimum threshold of copy number for each element) for AnnoSINE_v2; “-t” (target site duplications (TSD) mismatch tolerance) and “-s” (TSD score cutoff) for SINE-Finder). TPR: true positive rate. FPR: false positive rate.

**Fig. 3.**
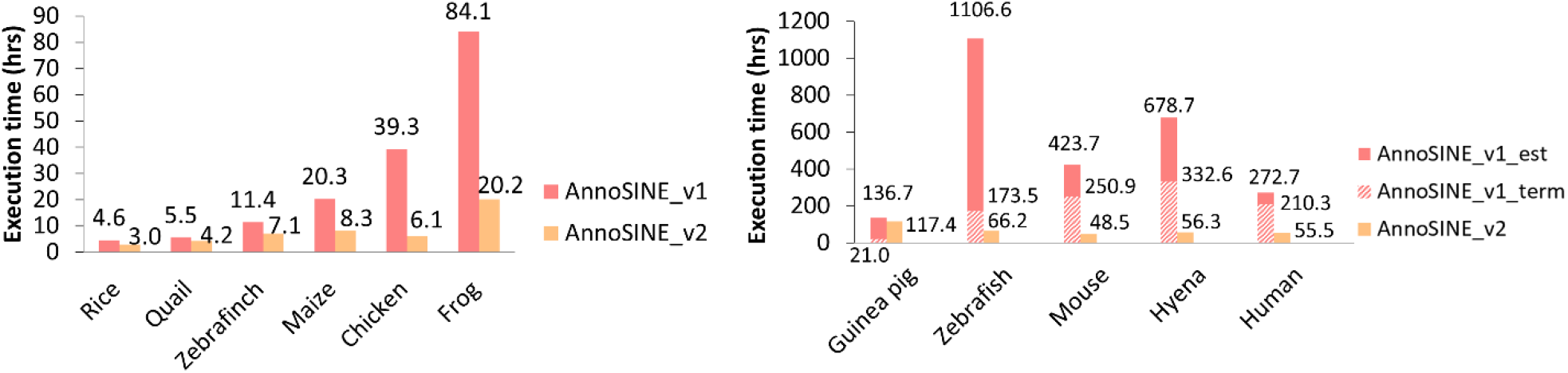
The execution time of AnnoSINE_v1 and AnnoSINE_v2 on plant and animal genomes. Executions of AnnoSINE_v1 on genomes shown on the right panel were interrupted due to overconsumption of the server memory. The recorded execution time until termination was labeled as “Annosine_v1_term”. The required times of AnnoSINE_v1 were estimated (“AnnoSINE_v1_est”) based on the progress where it was interrupted. Each benchmark was executed using 20 CPU cores (AMD EPYC 7763 2.45 GHz) and 512 GB of memory (DDR4 3200 MHz).

In addition to its proficiency in annotating SINEs in animal genomes, AnnoSINE_v2 has undergone several optimizations that significantly improve its computational performance. We benchmarked AnnoSINE_v2’s execution time with that of AnnoSINE_v1 across 11 plant and animal genomes. To enable AnnoSINE_v1 for animal genomes, we incorporated the animal pHMM database used in AnnoSINE_v2 while keeping all other components unchanged. In all experiments, AnnoSINE_v1 was run with one core, and AnnoSINE_v2 was run using 20 cores. For five of the tested genomes (guinea pig, zebrafish, mouse, hyena, and human), AnnoSINE_v1 could not be finished due to the out-of-memory error for its over-utilization of the 512 GB server memory. Consequently, we recorded the execution time until termination and estimated the execution time of AnnoSINE_v1 on these genomes based on the progress where it was terminated. Our results demonstrate that AnnoSINE_v2 completes the SINE annotation task in much less time compared to AnnoSINE_v1. Notably, it achieves over a 5-fold speed up in the chicken genome compared to AnnoSINE_v1 and demonstrates an acceleration of over 150 and 250 hours in the human and hyena genomes, respectively. To further investigate the speed improvements of AnnoSINE_v2, we applied AnnoSINE_v1 with one core and AnnoSINE_v2 with varying cores to analyze the chromosome 1 of the human genome. However, AnnoSINE_v1 was terminated prematurely due to overconsumption of server memory. The result shows that AnnoSINE_v2 achieves over an 8-fold speed improvement (over 110 hours) compared to AnnoSINE_v1 when using a single core (Fig. 4), which demonstrates the speed improvement coming from the algorithm optimization. In addition, we observed that AnnoSINE_v2 exhibits more than a 2-fold speed improvement with only 4 cores, while the rate of improvement gradually diminishes as more cores are utilized. These findings suggest that AnnoSINE_v2 is an efficient SINE annotation tool for both plant and animal genomes.

**Fig. 4.**
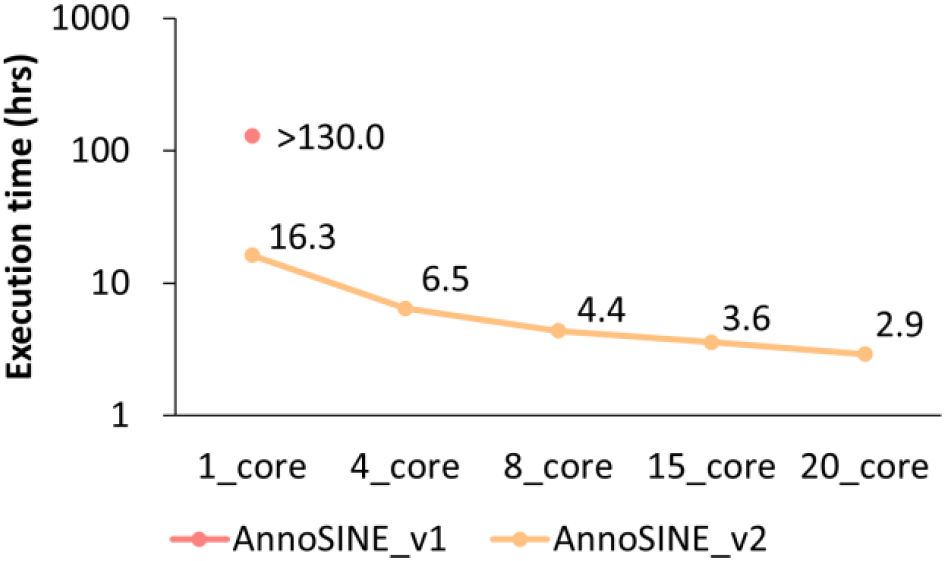
The execution time of AnnoSINE_v1 and AnnoINE_v2 on human chromosome 1. The y-axis is shown in log-scale. Execution of AnnoSINE_v1 on the test genome was terminated due to overconsumption of server memory, and the recorded time reflects the execution time until termination. Each benchmark was executed using Intel Xeon E5-2680 v4 (2.40 GHz and 128 GB of memory (DDR4 2400 MHz).

## Materials and Methods

### Genomes used in this study

Apple (*Malus domestica*, GCF_002114115.1), Arabidopsis (*Arabidopsis thaliana*, Col-CC, GCA_028009825.2), chicken (*Gallus domesticus*, galGal6, GCF_016700215.2), frog (*Rana temporaria*, GCF_905171775.1), guinea pig (*Cavia porcellus*, GCF_000151735.1), human (*Homo sapiens*, hg38, GCF_000001405.40), hyena (*Crocuta crocuta*, GCA_008692635.1), maize (*Zea mays*, B73v5, GCF_902167145.1), mouse (*Mus musculus*, GRCm39, GCF_000001635.27), quail (*Coturnix japonica*, GCF_001577835.2), rice (*Oryza sativa*, MSUv7, GCF_001433935.1), zebra finch (*Taeniopygia guttata*, taeGut2, GCF_003957565.2), and zebrafish (*Danio rerio*, danRer11, GCF_000002035.6) genomes were used in this study.

### Extending SINE annotations on animal genomes

Despite AnnoSINE’s effectiveness in annotating plant genomes, it has limited capability to annotate SINE elements in animal genomes due to the lack of animal-based pHMMs for the homology search function. To enhance the SINE annotation in animal genomes, we initially obtained 139 pHMMs from the DFAM database [6] with “SINE” and “animal” keywords in their annotations. With manual inspection, we determined that 21 of these pHMMs corresponded to non-SINE elements. Specifically, to determine whether downloaded pHMMs are SINE elements, we performed manual identification.

Specifically, we considered the pHMMs to be non-SINE elements if their “Description” and “Classification and Taxa” entries in the DFAM page did not contain the word “SINE”, any SINE-related descriptions or contain non-SINE descriptions. The remaining 118 pHMMs were then added to AnnoSINE_v2 to enhance the pHMM-based homology search. To distinguish with SINE annotations in plant genomes, we added the new parameter “-a” for this function, with “-a 0” retaining the original function for plant SINE element searches, “-a 1” for animal SINE elements, and “-a 2” that allows the identification of both plant and animal SINE elements.

### Optimizing SINE-Finder

SINE-Finder (v1.0.1) [5] has been incorporated into AnnoSINE (v1) for *de novo* SINE annotations. In our previous experiments, we observed that the SINE-Finder step often required a significant amount of time. Upon thorough examination, we found that the SINE-Finder program is highly sensitive to the input genome format that can vary from users. Thus, we employed Seqtk (https://github.com/lh3/seqtk) to convert the input genomes into the single-line fasta format as the input for the SINE-Finder function.

Furthermore, we used the chunking parameter available in SINE-Finder with a chunk size of 10kb to allow efficient processing of large genomes. These adaptations significantly reduced the processing time. For instance, this optimization resulted in a decrease in the processing time of the apple genome (591 Mb) from 30,000 seconds to 800 seconds.

### Optimized pHMM merging algorithm

The original pHMM merging function in AnnoSINE is designed to merge overlapping pHMM alignments, but it suffers from low efficiency due to the presence of a nested loop that recurrently traverses each alignment for the identification of overlaps. In order to enhance the efficiency of this function, we developed an optimized version that incorporates a sorting strategy to arrange the list of pHMM alignments. With the sorted list, the function examines overlaps and employs a dictionary to store the overlapping records, and the merging will be carried out after traversing the sorted list. Suppose n refers to the number of alignment records output by HMMER. The original strategy requires traversing the whole list for each alignment record (the nested loop) with n*n times of iteration. The sorting strategy allows the function to iterate through all items only once, resulting in a total of n iterations. By using the sorting (*O*(*nlogn*)) and a linear merging algorithm (O(*n*)), this new approach reduces the time from *O*(*n*^2^) to *O*(*nlogn*), leading to a significant improvement of speed.

### Versatile strategies for analyzing large genomes

BLAST was employed to identify the copy number of SINE candidates in AnnoSINE. However, BLAST is inefficient in processing many input sequences and usually generates TB-sized results that hinder the parsing efficiency. To address these issues, we replaced BLAST with Minimap2 [9], a versatile and efficient aligner specifically designed for high-throughput alignment tasks. Notably, Minimap2 allows secondary alignments by adjusting the parameter “-N”. In addition, its capability to handle structural variation is crucial for accurately aligning highly divergent SINE copies. Moreover, Minimap2 improves alignment efficiency and reduces file complexity compared to BLAST. Thus, using Minimap2 can enhance the overall performance and accuracy of the AnnoSINE program. To allow this Minimap2 conversion, we first changed the BLAST output from the original alignment format 1 to the tabular format 6 and adapted AnnoSINE to process the new BLAST format. Then, we developed a custom script to convert the Minimap2 PAF format to the BLAST format 6 to maintain compatibility.

To accelerate the format conversion in large PAF files, we implement a chunking strategy to split the PAF file into 100k lines per chunk and process chunks with user-specified multi-threads. Processing large PAF files will consume a significant amount of memory, which may exceed the server’s limit. To avoid such errors, we implemented a strategy in the conversion script to store BLAST format 6 outputs into files with approximately 100 million alignments per file, which is equivalent to ∼10 GB per file and only leaves a small memory footprint.

In our tests, the Minimap2 component still could drain the system memory for large input genomes and cause program interruption. To address this issue, we implemented a function to automatically switch Minimap2 to the memory-efficient mode (the “-K 1M” parameter) if the SINE candidate file exceeds the preset size (default:10 Mb). These improvements allow the processing of large genomes while remaining efficient.

### Speedup by using multiple CPU threads

We implemented parallel computation in AnnoSINE_v2 where possible. Firstly, we introduced a parallel searching strategy for the TSD searching function. This approach divides the input sequence into n parts (n being the number of threads set by the user, default is 8) and performs parallel TSD searches on these divided sequences.

Additionally, we passed the multithreading option to other tools, such as Minimap2 and HMMER, within the program. These enhancements contribute to an overall faster and more efficient execution of the program. We also adapted versatile codes in the program to allow its execution outside of the program directory, allowing easy installation and utility.

### Benchmarking program performances

We benchmarked the annotation performance of SINE-Finder and AnnoSINE_v2 in zebrafish, human, and mouse genomes in comparison to their standard annotations obtained from the UCSC genome browser [6]. The optimized SINE-Finder was used in these benchmarks. To remove redundant sequences in SINE-Finder’s predictions, the “cleanup_nested.pl” script from the EDTA package [10] was used with parameters “-cov 0.95 -minlen 50 -miniden 80”, which identified and removed redundant copies with coverage ≥ 95%, identity ≥ 80%, and alignment length ≥ 50 bp. Since AnnoSINE_v1 was designed for plant genomes, and its executions were interrupted on the three genomes due to out-of-memory error, we could not benchmark AnnoSINE_v1.

AnnoSINE_v2 was executed on these animal genomes using the “-a 2” parameter, which used both plant and animal pHMMs for the homology search. The resulting SINE predictions were purged using LINE and LTR retrotransposon libraries from respective genomes generated by EDTA (v2.2.0). The purging was achieved using RepeatMasker (v4.0.6) (http://www.repeatmasker.org) and the “cleanup_tandem.pl” script from the EDTA package with parameters “-Nscreen 0 -nr 0.8 -minlen 50 -cleanN 1 -cleanT 1 -trf 0”. Further, the resulting SINE libraries were used to annotate genomes with RepeatMasker and parameters “-q -no_is -nolow -div 40 -cutoff 225”, which allows for identifying SINE-related sequences with up to 40% divergence. Finally, the whole-genome RepeatMasker annotation was compared against the UCSC standard annotation using the “lib-test.pl” script from the EDTA package.

To benchmark the computational performance of AnnoSINE_v1 and AnnoSINE_v2, we executed these programs on 11 plant and animal genomes using 20 CPU cores (AMD EPYC 7763 2.45 GHz) and 512 GB of memory (DDR4 3200 MHz). AnnoSINE_v1 could not finish its execution in five animal genomes due to their large genome size and the abundance of SINE candidates. Since the program was interrupted during the BLAST step, we were able to estimate the required time of execution based on the number of processed SINE candidates relative to the total number of candidates and the time past before the interruption.

## Conclusion and Discussion

In this work, we presented AnnoSINE_v2, a new version of AnnoSINE for efficient SINE annotation in plant and animal genomes. By incorporating animal pHMMs, AnnoSINE_v2 can identify not only plant SINEs but also animal SINEs. In addition, AnnoSINE_v2 is more computationally efficient than AnnoSINE_v1 through algorithmic optimization and the utilization of multithreading. Through benchmarking, we also exposed opportunities to further improve the sensitivity of SINE annotation on certain animal genomes (e.g. mouse). Thus, for the future work, we aim to improve the sensitivity of our tool while keeping the high specificity. For example, applying AnnoSINE_v2 in structural mode to identify novel SINEs and incorporating new pHMMs with manual correction can be considered.

## Declarations

## Ethics approval and consent to participate

Not applicable.

## Consent for publication

Not applicable.

## Availability of data and materials

AnnoSINE_v2 is freely available under MIT license on Conda and GitHub: https://github.com/liaoherui/AnnoSINE_v2.

## Authors’ contributions

SO and YS designed the study. HL and SO developed the code. SO and HL collected and analyzed the data. HL, SO, and YS wrote and revised the paper.

## Acknowledgements

The software provided in this work was “as-is” and does not constitute a guarantee or warranty by OSU. We would like to thank ChatGPT for helping with drafting and improving some of the scripts used in this study.

## Abbreviations

SINEs: Short interspersed nuclear elements
pHMMs: profile hidden Markov models
TPR: True positive rate
FPR: False positive rate
TSD: Target site duplications

